# Translational profiling reveals novel gene expression changes in the direct and indirect pathways in a mouse model of LDOPA-induced dyskinesia

**DOI:** 10.1101/2024.06.10.598251

**Authors:** Sabika Jafri, Mahdi Ghani, Natalie Stickle, Carl Virtanen, Lili-Naz Hazrati, Naomi P. Visanji

**Affiliations:** Krembil Brain Institute, University Health Network, Toronto, ON, Canada; Tanz Centre for Research in Neurodegenerative Disease, University of Toronto, Krembil Discovery Tower, 60 Leonard Ave, Toronto, ON M5T 0S8, Canada; Department of Laboratory Medicine and Pathobiology, University of Toronto, Toronto, ON, Canada; University Health Network Microarray Centre, Toronto Medical Discovery Tower, 101 College Street, Rm 9-301, Toronto, Ontario M5G 1L7, Canada; Department of Pathology, McGill University, 3801 University Avenue, Montreal, Qc, H3A 2B4, Canada

**Keywords:** Parkinson’s disease, levodopa, dyskinesia, gene expression

## Abstract

The molecular mechanisms underlying L-dihydroxyphenylalanine (LDOPA) induced dyskinesia in Parkinson’s disease are poorly understood. Here we employ two transgenic mouse lines, combining translating ribosomal affinity purification (TRAP) with bacterial artificial chromosome expression (Bac), to selectively isolate RNA from either DRD1A expressing striatonigral, or DRD2 expressing striatopallidal medium spiny neurons (MSNs) of the direct and indirect pathways respectively, to study changes in translational gene expression following repeated LDOPA treatment. 6-OHDA lesioned DRD1A and DRD2 BacTRAP mice were treated with either saline or LDOPA bi-daily for 21 days over which time they developed abnormal involuntary movements reminiscent of dyskinesia. On day 22, all animals received LDOPA 40 minutes prior to sacrifice. The striatum of the lesioned hemisphere was dissected and subject to TRAP. Extracted ribosomal RNA was amplified, purified and gene expression was quantified using microarray. 195 significantly varying transcripts were identified among the 4 treatment groups. Pathway analysis revealed an overrepresentation of calcium signaling and long-term potentiation in the DRD1A expressing MSNs of the direct pathway, with significant involvement of long-term depression in the DRD2 expressing MSNs of the indirect pathway following chronic treatment with LDOPA. Several MAPK associated genes (*NR4A1*, *GADD45G*, *STMN1*, *FOS* and *DUSP1*) differentiated the direct and indirect pathways following both acute and chronic LDOPA treatment. However, the MAPK pathway activator *PAK1* was downregulated in the indirect pathway and upregulated in the direct pathway, strongly suggesting a role for *PAK1* in regulating the opposing effects of LDOPA on these two pathways in dyskinesia. Future studies will assess the potential of targeting these genes and pathways to prevent the development of LDOPA-induced dyskinesia.

## Introduction

Since its first use in the early 1960’s, dopamine replacement, using the dopamine precursor L-dihydroxyphenylalanine (LDOPA), has remained the most effective therapy for the motor symptoms of Parkinson’s disease (PD). However, following long-term treatment with LDOPA, >90% of PD patients develop highly debilitating abnormal involuntary movements termed LDOPA induced dyskinesia (LID) (A. Fabbrini & Guerra, 2021; G. Fabbrini et al., 2007; Obeso et al., 2000; Rascol, 2000). In 2014 a Priority Setting Partnership, commissioned by Parkinson’s UK, identified LID as 3rd of 96 unmet needs in PD (Deane et al., 2015). The socioeconomic impact of LID places an enormous burden on the patient, caregiver and healthcare system (Dodel et al., 2001), however, as the pathogenesis of LID is poorly understood, treatment options are very limited (AlShimemeri et al., 2020). Once developed, LID is virtually impossible to reduce or reverse and sensitivity to developing dyskinesia on re-exposure persists even after long periods of treatment discontinuation.

Behavioral sensitization to repeated LDOPA treatment is also seen in animal models of PD (Angela Cenci & Lundblad, 2007; Bezard et al., 2001; Jenner, 2008; Lundblad et al., 2002, 2004, 2005). In the 6-hydroxydopamine (6-OHDA) lesioned rodent, sensitization to repeated LDOPA treatment leads to abnormal involuntary movements (AIMs) reminiscent of LID. These rodent behaviors have been shown to have a similar pharmacology to LID in humans (Lundblad et al., 2005). Extensive studies in these animal models have revealed that dopamine can directly modulate the functioning of the striatum and that the pathophysiology of LID involves a wide range of changes in the basal ganglia which have been characterized using a variety of methods. These studies have found that both presynaptic and postsynaptic mechanisms are important in governing the manifestation of LID, with changes in the regulation of several genes as well as alterations in the electrophysiological activities of striatal circuits and synaptic plasticity (Borgkvist et al., 2018; Huot et al., 2013; Hutny et al., 2021; Mosharov et al., 2015; Spigolon & Fisone, 2018; Surmeier et al., 2014).

Overstimulation of postsynaptic dopamine receptors located on GABAergic medium spiny neurons (MSNs) in the dorsal striatum of the basal ganglia have been shown to be the site of the initial generation of LID (Fisone & Bezard, 2011). The MSNs form two distinct projection pathways, exerting opposing effects on motor activity: the direct DRD1A dopamine receptor expressing pathway projects to the substantia nigra and promotes locomotion, whereas the indirect DRD2 dopamine receptor expressing pathway projects to the pallidum and inhibits locomotion (Albin et al., 1989; Alexander & Crutcher, 1990). To fully understand how dopaminergic stimulation at the level of the MSNs leads to the development of LID, it is vital to study the molecular responses of these two distinct populations of MSNs to LDOPA in isolation (Cenci et al., 2018; Ryan et al., 2018). However, striatal MSNs are both anatomically intermixed and morphologically indistinguishable (Calabresi et al., 2014), thus an inability to adequately discriminate the MSNs of the direct and indirect pathways has been a significant roadblock to the full understanding of the development of LID. In the present study, and as we have previously, we have made use of two transgenic mouse lines combining translating ribosomal affinity purification (TRAP) with bacterial artificial chromosome (Bac) technology to purify RNA from either DRD1A expressing striatonigral MSNs or DRD2 expressing striatopallidal MSNs (Dougherty, 2017; Heiman et al., 2008, 2014; Visanji et al., 2015). Briefly, these mice express an enhanced green fluorescent protein (EGFP) tagged L10a ribosomal subunit under the control of either the DRD1A or DRD2 receptor promoter, resulting in the expression of EGFP-L10a in the MSNs of either the direct or indirect pathway, respectively. The EGFP-L10a subunits of the MSNs of both the direct and indirect pathways can be purified along with the associated mRNA which can then be profiled using microarray to reveal differences in translational RNA expression in these two distinct cell populations. Indeed, translational profiling in these two mouse lines has previously identified >70 transcripts enriched in the indirect pathway and >150 transcripts enriched in the direct pathway, as well as validating the differential expression of known markers that distinguish these two cell populations (Heiman et al., 2008).

Here, we employ a widely used model of LID in 6-OHDA lesioned rodents, using two transgenic mouse lines to selectively identify changes in translational gene expression induced by chronic LDOPA in either DRD1A expressing striatonigral (direct pathway) or DRD2 expressing striatopallidal (indirect pathway) MSNs and reveal mechanisms underlying LID that might aid the development of preventative strategies for one of the major challenges in the treatment of PD.

## Methods

### Transgenic mice

All animal use was in accordance with approved local institution protocol and the regulations defined by the Canadian Council on Animal Care. Two Bac transgenic mouse lines were obtained from The Rockefeller University, New York, NY, USA. Both transgenic mouse lines express an EGFP-L10a fusion protein either under the control of the DRD1A (line CP73) or DRD2 (line CP101) promoter. For a full description of the mice, please refer to (Doyle et al., 2008). Both lines were on a C57BL/6J/Swiss-Webster background and were maintained as transheterozygous.

### 6-hydroxydopamine (6-OHDA) lesioning of the median forebrain bundle (MFB)

All animals were lesioned at 35 days of age. 30 minutes prior to lesioning, animals received desipramine (25mg/kg) and pargyline (5mg/kg) intraperitoneally (i.p.) (Both Sigma Aldrich). Under anaesthesia (isoflurane), 6-OHDA (3ug in 0.6ul) (Sigma Aldrich) was infused unilaterally into the medial forebrain bundle (MFB) at a flow rate of 0.2ul/min at the following coordinates from Bregma: AP -1.2, ML -1.1, DV -5.0 mm, according to the mouse brain atlas (George Paxinos & Keith B.J. Franklin, 2013). The needle was left in situ for 5 minutes before being retracted. Animals had a 14-day recovery period during which time they were administered 3ml/day lactated ringers containing 5% dextrose and kept on heat pads until they were able to maintain a stable body weight. 14 days post-lesion, all animals were assessed for rotational bias by measuring spontaneous full rotations contraversive and ipsiversive to the lesioned hemisphere over a 10 minute period in glass cylinders. Post-mortem, lesion efficiency was assessed by calculating nigral tyrosine hydroxylase (TH) immunoreactive cell loss. Following fixation in 4% paraformaldehyde, blocks encompassing the entire midbrain were embedded in paraffin and 5 micron-thick serial sections were taken from -3.08mm and -3.28mm relative to bregma, according to the mouse brain atlas (George Paxinos & Keith B.J. Franklin, 2013). Briefly, endogenous peroxidase was blocked with 3% hydrogen peroxide, antigen retrieval was done using 10 mM citrate buffer at pH 6.0, and sections were stained overnight with rabbit polyclonal antibody to TH (Novus Biologicals) at 1/1500 dilution. The staining was finished with Vector’s Peroxidase ImmPRESS detection system, the color was developed by 3,31-Diaminobenzidine, and sections were counterstained with Mayer’s hematoxylin prior to being coverslipped. TH positive cells were counted manually and defined as those with a brown cell membrane and distinct lighter rounded cell body.

### LDOPA treatment and abnormal involuntary movement (AIMS) analysis

All animals were administered either vehicle, or LDOPA methyl ester/benserazide (6/1.5mg/kg) (Sigma Aldrich) i.p. twice daily (>6 hours apart) for 21 days. On days 1, 7, 14 and 21 of treatment, animals were assessed for abnormal involuntary movements (AIMs). Immediately post treatment, animals were placed in single cages and their behaviour was observed for 1 minute every 20 minutes for a period of 3 hours. The AIMs scale, as described in detail by Cenci and Lundblad., 2007, was used to assess the level of dyskinesia (Angela Cenci & Lundblad, 2007). Briefly, there were 3 categories of scored AIMs: axial, limb, and orofacial. Each category was rated on a scale of 1-4 based on the maximum severity of behaviour observed in each 1 minute period. Please refer to Supplemental Table 1 for a full description of each AIMs category and rating.

### Translating ribosome affinity purification (TRAP)

On day 22, all animals were sacrificed 40 minutes post treatment with LDOPA/benserazide (6/1.5mg/kg). Mice were killed by cervical dislocation and the striatum of the lesioned hemisphere was quickly dissected on ice. Dissected striata were individually homogenised, using a motor driven Teflon-glass homogeniser, in ice cold lysis buffer (20mM HEPES KOH, 5mM MgCl2, 150mM KCL, 0.5mM DTT, 100ug cyclohexaminde, 40U/ml Rnasin and protease inhibitor cocktail, pH 7.4). A post-nuclear supernatant was prepared by centrifugation at 2,000 x g at 4°C for 10 min to which 1% NP-40 (EMD Millipore, Billerica, Massachusetts) and 30mM 1,2-diheptanoyl-sn-glycero-3-phosphocholine (DHPD, Avanti Polar Lipids, Alabaster, AL) was added. Finally, a post mitochondrial supernatant was prepared by centrifugation at 20,000 x g at 4°C for 10 min. Striatal homogenates were immunoprecipitated by end-over mixing for 30 min at 4°C with Dynal protein G magnetic beads (Invitrogen, Carlsbad, CA) pre-coated with 50ug each of two anti-GFP antibodies (clones 19C8 and 19F7, Memorial Sloane Kettering, NYC, NY). Following immunoprecipitation, beads were collected using a magnetic rack and washed four times with an ice-cold high salt buffer (20mM HEPES-KOH, 5mM MgCl_2_, 350mM KCL, 0.5mM DTT, 1% NP-40, 100ug cyclohexaminde, pH 7.4). RNA was extracted from the beads and purified according to the manufacturer’s instructions using an Absolutely RNA nanoprep kit (Stratagene, La Jolla, CA) with in column DNA digestion and frozen prior to gene expression profiling being performed. The quantity of RNA was determined using a Nanodrop 1000 spectrophotometer (Wilmington, DE) and the quality was determined using an Agilent 2100 Bioanalyzer (Foster City, CA).

### Microarray

1ng of extracted RNA from each striatal sample (Stratagene/Agilent) was amplified using the WT-Ovation Pico RNA amplification System Version 1.0 (Nugen) and cDNA was run on a bioanalyzer (Agilent) for quality control. All samples had a RIN number of >6.3. 3.5ug cDNA was biotin labeled (Nugen Illumina) 1.5ug of cDNA was hybridized to the Illumina Mouse WG-6 V2.0 BeadChip containing 45821 probes. BeadChips were incubated at 48°C with a rotation speed of 5 for 18.0 hrs of hybridization. BeadChips were then washed and stained as per Illumina protocol and scanned on the iScan (Illumina). Data files were quantified in GenomeStudio Version 2010.2 (Illumina). All samples passed Illumina’s sample dependent and independent quality control metrics. Data were further checked for overall quality using R (v2.14.1) with the Bioconductor framework and the LUMI package installed. There were no discernible outliers.

Data was imported into Genespring v12v12.0 (Agilent) for analysis and normalized with a quantile normalization followed by a median centering. All data analysis and visualization were performed on log2 transformed data. There were 4 groups overall split into 2 factors: MSN type (Drd1 EGFP-L10a expressing, DRD2 EGFP-L10a expressing) and LDOPA treatment (acute or chronic). Data was first filtered to remove the confounding effect probes that show no signal may have on subsequent analysis. Only probes that were in the upper 80th percentile of the distribution of intensities in 100% of any of the 1 of 4 above groups were allowed to pass through this filtering. The final set for analysis contained 32362 probes. To find those genes that statistically varied with the greatest confidence between sample groups, a 1-way ANOVA with a Benjamini-Hochberg False Discovery Rate (q<0.05) was used. In order to look for specific comparisons of interest a post-hoc Tukey’s Honest Significant Difference (HSD) test was used after the ANOVA.

### Pathway analysis

Enrichr (http://amp.pharm.mssm.edu/Enrichr/) was applied to identify significant Kyoto Encyclopedia of Genes and Genomes (KEGG) defined pathways associated with the set of differentially expressed genes in each experimental group.

### Quantitative Real-Time RT-PCR

Remaining cDNA from the amplified product used for microarray was used for RT-PCR. PCR was performed using Promega GoTaq qPCR mastermix according to the manufacturer’s instructions (Promega, Madison, WI). Each reaction comprised 1 ng cDNA, 12.5 ul GoTaq and 0.2 uM final concentration of each primer. Cycling and detection were carried out using an Applied Biosystems 7500 Real Time PCR System and data quantified using Sequence Detection Software Version 1.4 (Applied Biosystems, Carlsbad, CA). PCR was performed for a total of 40 cycles (95° for 15 sec, 60° for 60 sec,) followed by a dissociation stage. Each sample was assayed in duplicate. All data were normalised to Actin and quantification was carried out via the absolute method using standard curves generated from pooled cDNA representative of each sample to be analysed. Please refer to Supplemental Table 2 for a complete list of all primers used in the present study.

### Statistical analysis

All data, other than the microarray data analysis described above, were analysed using Prism 5 (GraphPad, La Jolla, CA).

## Results

### Characterisation of 6-OHDA lesion of the nigrostriatal pathway and abnormal involuntary movements following chronic treatment with LDOPA in Drd1 and Drd2 EGFP-L10a Bac mice

Successful lesion of the nigrostriatal pathway was confirmed both behaviourally and histologically. Thus, 14 days post-lesion, all animals were assessed for rotational bias by measuring spontaneous full rotations contraversive and ipsiversive to the lesioned hemisphere over a 10 minute period in glass cylinders. All animals included in the study displayed a ≥90% ipsiversive rotational bias (see Supplemental Table 3). Post-mortem, lesion efficiency was assessed by calculating nigral tyrosine hydroxylase (TH) immunoreactive cell loss. All animals included in the study had a ≥90% loss of tyrosine hydroxylase immunoreactive cells in the lesioned hemisphere as compared to the intact hemisphere (see Supplemental Table 3).

Following 21 days treatment with LDOPA, both DRD1A and DRD2 animals exhibited significantly higher levels of AIMs compared to animals in the acute treatment group who had been treated with saline for 21 days (Supplementary Figure 1). There was no significant difference in the amplitude of peak dose AIMs between Drd1 and Drd2 EGFP-L10a Bac mice.

### Striatal gene expression analysis following acute or chronic treatment with LDOPA in DRD1A and DRD2 EGFP-L10a Bac mice

To find the genes that statistically varied with the greatest confidence between sample groups, a 1-way ANOVA with a Benjamini-Hochberg False Discovery Rate (q<0.05) was used. In total, 207 probes representing 195 unique genes or transcripts were found with this test (Dataset 1). The results are clustered in Figure 1 for visualization purposes. Importantly, among the genes found, we were able to replicate known changes in the expression of *PDYN*, *HTR2A*, and *CREB1* following chronic LDOPA (Dataset 1), validating the use of our experimental paradigm to identify differences in the molecular responses of the direct and indirect pathway to chronic LDOPA.

**Figure 1.**
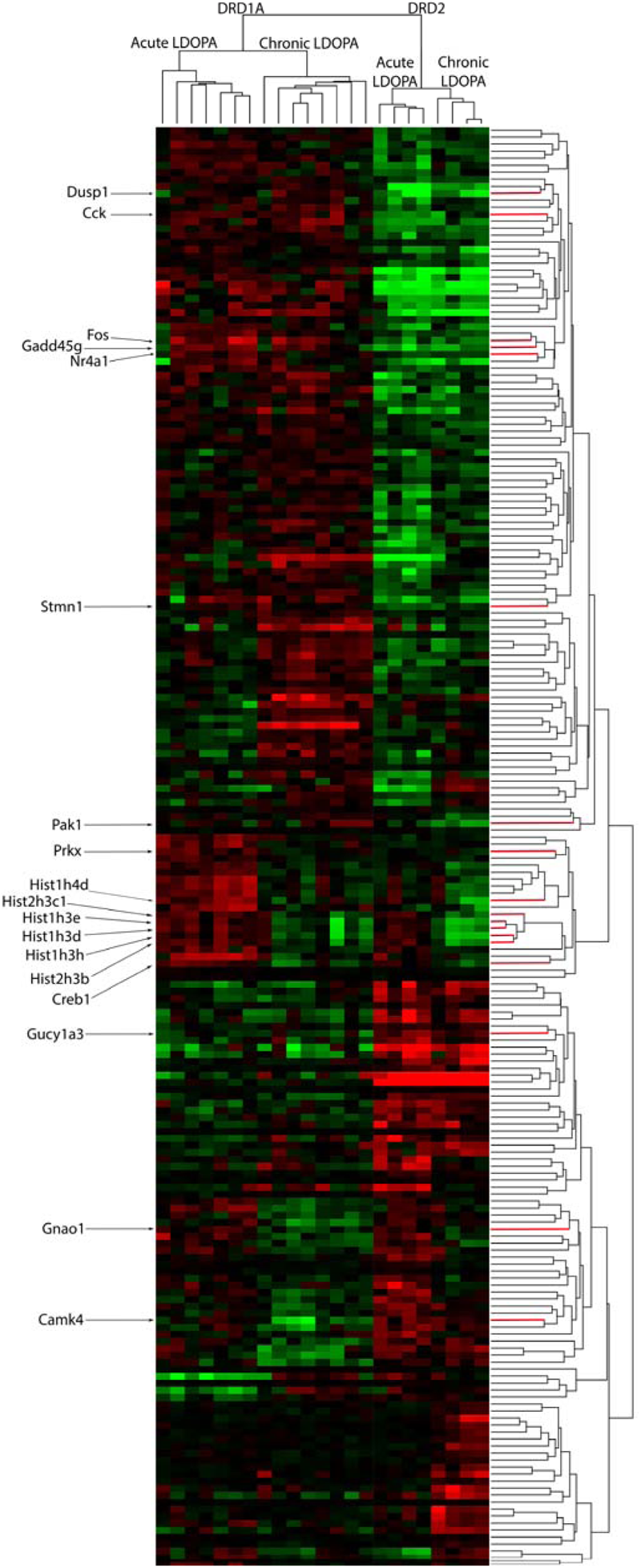
Heat-map of supervised clusters of gene expression changes in Drd1a and Drd2 EGFP-L10a expressing MSNs following acute or chronic LDOPA in a mouse model of PD. A 2-way hierarchical clustering of the 207 significantly varying probes, as determined through a Benjamini-Hochberg FDR corrected ANOVA (q<0.05), is presented. Color indicates direction of and magnitude of log2 fold-change (red, increased; green, decreased; black, no significant change). The dendrogram above the heatmap illustrates the similarity of samples within each group, the dendrogram to the right shows the hierarchical clustering based on similarity between the expression of genes. Specific genes of interest are named on the left.

A post-hoc Tukey’s HSD test was applied to the 207 probes of interest to perform select pairwise comparisons. Probes altered between DRD1A and DRD2 MSNs in animals treated with acute LDOPA are found in Dataset 2. Probes altered in DRD1A MSNs following acute vs chronic LDOPA treatment are found in Dataset 3. Probes altered in DRD2 MSNs following acute vs chronic LDOPA treatment are found in Dataset 4. Probes altered between DRD1A and DRD2 MSNs after chronic LDOPA treatment are found in Dataset 5. Venn analysis was used to generate lists of gene expression changes both common to and exclusive to Drd1 and DRD2 EGFP-L10a expressing MSNs following acute and chronic LDOPA treatment (Figure 2). The largest dissimilarity was between the DRD1A and DRD2 EGFP-L10a expressing MSNs, followed by animals treated with acute versus chronic LDOPA. Thus, 129 genes were differentially expressed when comparing DRD1A and DRD2 EGFP-L10a Bac mice treated with acute LDOPA, 67 of which remained differentially expressed between DRD1A and DRD2 EGFP-L10a Bac mice treated with chronic LDOPA (Figure 2). In contrast, in DRD1A EGFP-L10a Bac mice 56 genes were differentially expressed when comparing animals treated with acute versus chronic LDOPA, and in DRD2 EGFP-L10a Bac mice 80 genes were differentially expressed when comparing animals treated with acute versus chronic LDOPA (Figure 2). Finally, 115 genes were differentially expressed when comparing DRD1A and DRD2 EGFP-L10a Bac mice treated with chronic LDOPA, 48 of which were not differentially expressed when comparing DRD1A and DRD2 EGFP-L10a Bac mice treated with acute LDOPA (Figure 2).

**Figure 2.**
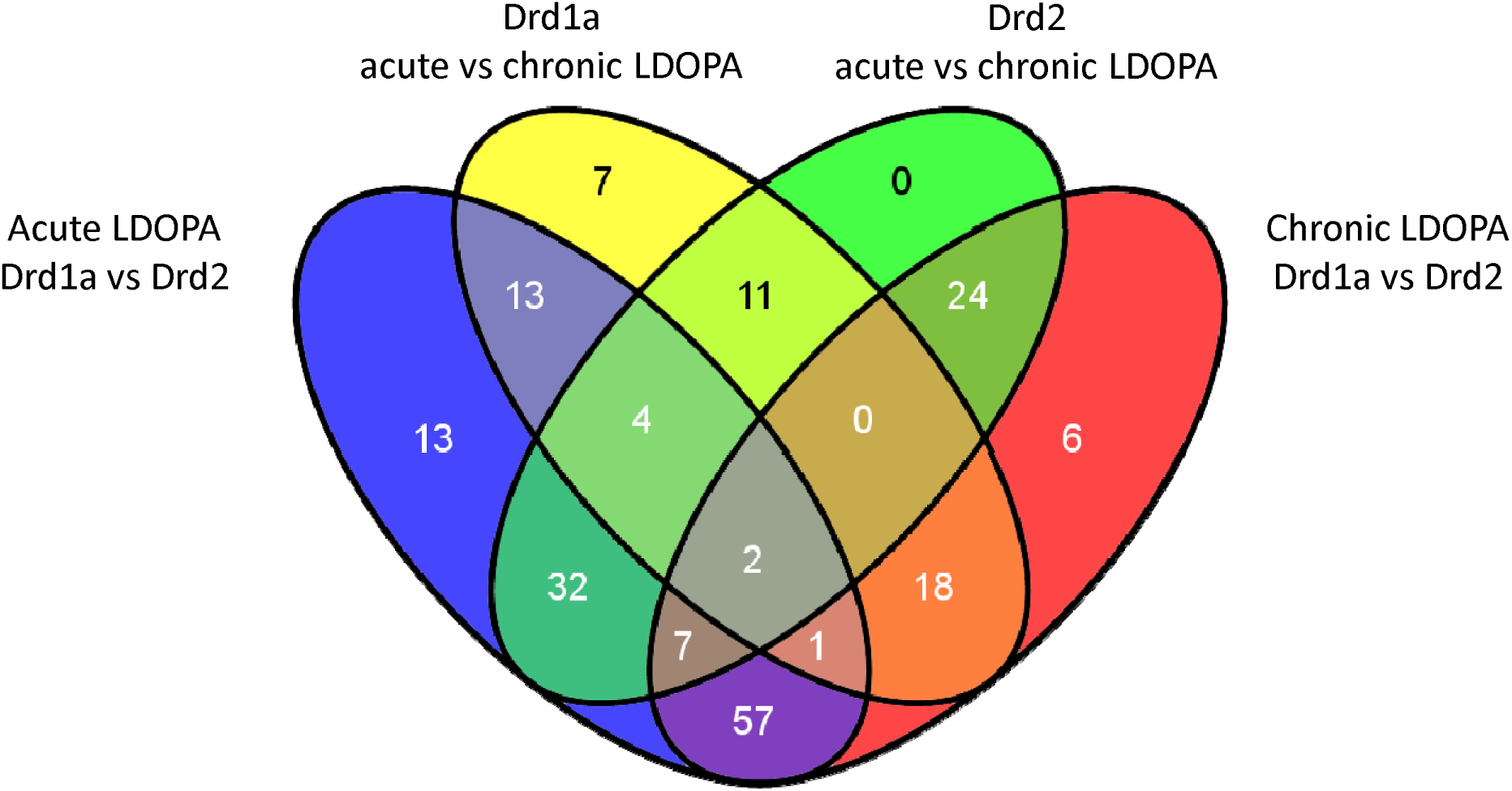
Overlap of differentially expressed genes in Drd1a and Drd2 EGFP-L10a expressing MSNs following acute and chronic LDOPA in a mouse model of PD. Venn diagram showing the total number of differentially expressed genes and their overlap between the four experimental groups for statistically significant changes (Benjamini-Hochberg FDR corrected ANOVA (q<0.05). Acute LDOPA Drd1a vs Drd2: Genes altered between Drd1a and Drd2 MSNs in animals treated with acute LDOPA. Drd1a acute vs chronic LDOPA: Genes altered in Drd1a MSNs following acute vs chronic LDOPA treatment. Drd2 acute vs chronic LDOPA: Genes altered in Drd2 MSNs following acute vs chronic LDOPA treatment. Chronic LDOPA Drd1a vs Drd2: Genes altered between Drd1a and Drd2 MSNs after chronic LDOPA treatment.

### Pathways altered in DRD1A and DRD2 EGFP-L10a expressing MSNs following acute and chronic treatment with LDOPA

The 195 unique genes that were differentially expressed among the four experimental groups were subjected to Enrichr analysis, which revealed several pathways differentially altered in DRD1A and DRD2 EGFP-L10a expressing MSNs following acute and chronic LDOPA as well as some common to both pathways (Table 1).

**Table 1.** KEGG defined pathways and their associated deregulated genes in Drd1a and Drd2 expressing EGFP-L10a MSNs following acute or chronic LDOPA in a mouse model of PD. Enrichr (http://amp.pharm.mssm.edu/Enrichr/) was applied to identify significant KEGG defined pathways associated with the sets of differentially expressed genes in Drd1a and Drd2 expressing MSNs following acute or chronic LDOPA. A: KEGG defined pathways associated with each of the four comparisons of interest. B: Table of differentially expressed genes associated with each KEGG defined pathway.

| Acute LDOPA Drd1a vs Drd2 |  |  |  |  |
| --- | --- | --- | --- | --- |
| KEGG Term | P-value | Zscore | Combined Score | Genes |
| HS04010_MAPK_SIGNALING_PATHWAY | 0.0001 | -2.16 | 21.42 | PRKX;GADD45G;DUSP1;DUSP6;STMN1;NR4A1;FOS;RASGRP4;RASGRP2 |
| HS04742_TASTE_TRANSDUCTION | 0.0066 | -1.96 | 9.85 | PRKX;GNL13;PDE1A |
| HS04740_OLFACTORY_TRANSDUCTION | 0.0217 | -1.70 | 6.53 | PRKX;GNAL |
| HS04540_GAP_JUNCTION | 0.0322 | -1.80 | 6.17 | PRKX;DRD2;GUCY1A3 |
| Drd1a acute vs chronic LDOPA |  |  |  |  |
| KEGG Term | P-value | Zscore | Combined Score | Genes |
| HS04916_MELANOGENESIS | 0.0002 | -1.97 | 17.11 | PRKX;DVL3;GNAO1;CREB1 |
| HS04020_CALCIIUM_SIGNALING_PATHWAY | 0.0012 | -1.91 | 12.77 | PRKX;CAMK4;GNAL;HTR2A |
| HS04740_OLFACTORY_TRANSDUCTION | 0.0039 | -1.70 | 9.46 | PRKX;GNAL |
| HS04720_LONG_TERM_POTENTIATION | 0.0171 | -1.70 | 6.90 | PRKX;CAMK4 |
| HS04540_GAP_JUNCTION | 0.0326 | -1.74 | 5.95 | PRKX;HTR2A |
| Drd2 acute vs chronic LDOPA |  |  |  |  |
| KEGG Term | P-value | Zscore | Combined Score | Genes |
| HS04510_FOCAL_ADHESION | 0.0082 | -2.17 | 10.43 | FARP2;MYL9;ITGA9;PAK1 |
| HS05120_EPITHELIAL_CELL_SIGNALING_IN_HELICOBACTER_PYLORI_INFECTION | 0.0325 | -1.79 | 6.13 | PAK1;ATP6V0A2 |
| HS04730_LONG_TERM_DEPRESSION | 0.0396 | -1.81 | 5.85 | GUCY1A3;GNAO1 |
| Chronic LDOPA Drd1a vs Drd2 |  |  |  |  |
| KEGG Term | P-value | Zscore | Combined Score | Genes |
| HS04010_MAPK_SIGNALING_PATHWAY | 0.0004 | -2.16 | 16.97 | GADD45G;DUSP1;STMN1;NR4A1;FOS;PAK1;RASGRP2 |
| HS04540_GAP_JUNCTION | 0.0174 | -1.91 | 7.75 | DRD2;GUCY1A3;HTR2A |
| HS00190_OXIDATIVE_PHOSPHORYLATION | 0.0343 | -1.74 | 5.87 | COX5A;NDUFA3;ATP6V0A2 |

The most significantly implicated pathway was the mitogen-activated protein kinase (MAPK) cascade (KEGG # HSA04010). Specifically, the significant MAPK pathway-associated genes *NR4A1*, *GADD45G*, *STMN1*, *FOS*, and *DUSP1*, were upregulated in the DRD1A expressing MSNs compared to DRD2 EGFP-L10a expressing MSNs in animals receiving both acute LDOPA (Enrichr p value 0.0001) and chronic LDOPA (Enrichr p value 0.0004). Likewise, the Gap junction pathway (KEGG # HSA04540) was also found to be associated when comparing DRD1A and DRD2 EGFP-L10a expressing MSNs following both acute and chronic LDOPA, with the former being driven by changes in the expression of *PRKX*, DRD2 and *GUCY1A3* (Enrichr p value 0.0332) and the latter by DRD2, *GUCY1A3* and *HTR2A* (Enrichr p value 0.0174) (Table 1).

DRD1A EGFP-L10a expressing MSNs had an overrepresentation of the long-term potentiation pathway (LTP) (KEGG # HSA04720; Enrichr p value 0.0171) after chronic treatment with LDOPA, whereas as DRD2 EGFP-L10a expressing MSNs had significant involvement of long-term depression (LTD) (KEGG # HSA04730; Enrichr p value 0.04) (Table 1). The LTP cascade was mainly driven by down-regulation of *PRKX* (KEGG #5613) and *CAMK4* (KEGG #814) in DRD1A EGFP-L10a expressing MSNs, with LTD in DRD2 EGFP-L10a expressing MSNs driven by upregulation of *GUCY1A3* (KEGG #2982) and downregulation of *GNAO1* (KEGG #2775).

### Validation of findings

To validate our findings, qRT-PCR was performed on 8 selected genes. First, the use of TRAP to successfully differentiate the MSNs of the direct and indirect pathways was confirmed by the selective enrichment of DRD1A and prodynorphin (*PDYN*) in RNA extracted from DRD1A EGFP-L10a Bac mice and of DRD2 and preproenkephalin 1 (*PENK1*) in RNA extracted from DRD2 EGFP-L10a Bac mice (Supplementary Figure 2).

Second, we chose four genes identified in the microarray as being differentially expressed across our 4 experimental groups to validate using qRT-PCR. Results of the qRT-PCR confirmed the microarray data. Thus, transcripts for *CAMK4*, *KCNK* and *SENP5* were all significantly upregulated and *TOM70A* was significantly downregulated in DRD2 EGFP-L10a expressing MSNs compared to DRD1A EGFP-L10a expressing MSNs following chronic treatment with LDOPA (Supplementary Figure 3).

## Discussion

Here, we describe translational gene expression changes in a mouse model of LID, employing a powerful methodology that allows for the selective study of DRD1A expressing striatonigral and DRD2 expressing striatopallidal projection MSNs. We have identified 195 unique genes altered in these two distinct neuronal subpopulations, implicated in the pathogenesis of LID, the majority of which are novel. Furthermore, pathway/GO analysis has revealed several differentially altered molecular pathways in which the gene expression changes were organized.

### Mitogen-activated protein kinase pathway and LID

The mitogen-activated protein kinase (MAPK) cascade, was the most significantly implicated GO pathway differentiating the DRD1A expressing (direct pathway) and DRD2 expressing (indirect pathway) MSNs, following both acute and chronic LDOPA treatment. The specific MAPK pathway related genes in our study included *DUSP1*, *STMN1*, *GADD45G*, *FOS* and *NR4A1*, all of which were upregulated in the MSNs of the direct pathway compared to the indirect pathway. This finding confirms previous observations for MAPK cascade activation in striatonigral DRD1A expressing MSNs, where an increase in *ERK* and *MSK1* phosphorylation was observed in dyskinetic mice in response to chronic LDOPA treatment (Santini et al., 2009) as well as a second study which, using the MAPK pathway gene *FOS*, combined optogenetics with the FosTRAP method and found that primarily direct pathway MSNs are activated during LID (Girasole et al., 2018).

DUSP1 is a phosphatase that dephosphorylates MAPK and suppresses activation of MAPK by oncogenic *RAS* (Wancket et al., 2012). A previous study found that *DUSP1* and *DUSP6* transcripts are upregulated in hemiparkinsonian mice subjected to LDOPA treatment (Pérez-Sen et al., 2019). Our data extend this finding and suggest this upregulation is specific to MSNs of the direct pathway. *STMN1* is a downstream effector of MAPK signalling via a miRNA-regulated mechanism and a potential oncogene in melanoma (Chen et al., 2013; Feng et al., 2017). *GADD45G* mediates activation of p38/JNK via MTK1/MEKK4 kinase (Balliet et al., 2003). *FOS* is one of the transcription factors regulated by MAPK signalling (Yokoyama et al., 2013). *NR4A1* encodes an intracellular transcription factor (Maxwell & Muscat, 2006). The involvement of these four genes in PD or LID has not been previously reported, although a previous study reported that FOSB and NURR1 (encoded by *NR4A2)* protein are elevated in dyskinetic rats (Steece-Collier et al., 2020). Elevated levels of FOSB have also been reported in dyskinetic non-human primates (Beck et al., 2019).

Notably, we observed that chronic treatment with LDOPA had opposing effects on the expression of PAK1, a known activator of the MAPK pathway and a breast cancer oncogene (Shrestha et al., 2012). Thus, *PAK1* gene expression was downregulated in the indirect pathway and upregulated in the direct pathway after chronic LDOPA treatment. As both the direct and indirect pathways play an important role in LID, this bidirectional alteration in gene expression make PAK1 particularly interesting. In addition to effects on MAPK, PAK1 is also a regulator of LTP via modulation of the actin cytoskeleton, with actin being heavily involved in receptor trafficking (Asrar et al., 2009). Altered subcellular trafficking of glutamatergic NMDA receptors has been implicated in LID, secondary to changes in the NR2A/NR2B subunit composition due to decreased NR2B levels resulting from deficient anchoring in the post-synaptic density (Cattabeni et al., 1999; Fiorentini et al., 2006). Given our observations future experiments might investigate the potential role of PAK1 in aberrant receptor trafficking in LID, with a view to potential therapeutic interventions.

### Cyclic AMP response-binding protein and LID

In our study, chronic LDOPA caused a downregulation of CREB1 RNA in both the direct and indirect pathways. Our observed implication of CREB1 in LID is consistent with another study using a transcriptomic approach, that reported that CREB1 was among the top upstream regulatory transcription factors implicated in rats with LID (Dyavar et al., 2020). In further support of a role for CREB1 in LID, Riluzole, known to reduce the activity of CREB1, has been shown to weaken LID in rats (Pagliaroli et al., 2019). It is widely understood that ERK and MAPK signalling regulate neuronal CREB transcription through phosphorylation (Koga et al., 2019) and previous studies have found that increased CREB phosphorylation is associated with LID as it is a downstream target of MSK1 (itself a target of ERK) (Azkona et al., 2014; Reyskens & Arthur, 2016). Administration of mGlu5R antagonists have been associated with reduced striatal levels of phosphorylated ERK and MSK1 as well as weakened dyskinesia in experimental PD (Spigolon & Fisone, 2018). Our findings provide further support for the role of CREB1 in both the direct and indirect pathways in LID and suggest that further studies investigating the potential of manipulating this pathway as possible treatment for LID are warranted.

### Histone cluster proteins and LID

Another target of ERK/MSK1 signalling is histone H3 and an increase in its phosphorylation has been reported in LID (Ciccarelli & Giustetto, 2014). In support of this, we found several histone cluster proteins (HIST1H3D, HIST1H4D, HIST1H3E, HIST1H3H, HIST2H3C1 AND HIST2H3B) to be downregulated in both the direct and indirect pathways following chronic treatment with LDOPA. Indeed, previous studies have found that long lasting cellular adaptations in animal models of LID include histone modifications through lysine acetylation, lysine and arginine methylation, serine and threonine phosphorylation, lysine ubiquitination and sumoylation, and the activation of histone kinases, histone acetyltransferases, histone deacetylases, and histone methyltransferases (reviewed in Cenci & Konradi, 2010). Such alterations affect the histone-DNA interaction as well as the ability of transcription factors to bind. We are not aware of any studies targeting modifications to histone cluster proteins as a potential therapeutic intervention in LID thus this would make a novel future direction.

### LTP, LTD and LID

Another major finding of our study was the involvement of calcium signalling with overlapping involvement of LTP in the direct pathway, and LTD in the indirect pathway in LID. Maladaptive plasticity in the striatal MSNs of the direct and indirect pathways in PD and LID has been previously described. Thus, in animal models of PD, corticostriatal LTP is lost but can be restored by treatment with LDOPA (Calabresi et al., 2007; Costa et al., 2012; Picconi et al., 2012). Interestingly, following induction of LTP, low-frequency stimulation can reverse the effect (depotentiation) but this ability is selectively lost in animals with LID (Picconi et al., 2012). These observations have led to the suggestion that striatal MSNs are able to manifest homeostatic adaptations in the number of excitatory corticostriatal synapses and intrinsic excitability in response to perturbations in dopamine signalling (Fieblinger et al., 2014).

In the present study, the significant implication of calcium signaling and LTP cascades in the direct pathway were mainly driven by down-regulation of PRKX and CAMK4 in chronic LDOPA treated animals compared to acute LDOPA treated animals. Both PRKX and CAMK4 are involved in LTP (Anderson & Kane, 1998), with calcium-dependent nuclear signalling via CAMK4 and CREB being involved in LTP-associated NMDA receptor post synaptic density-95 blockade (Bell et al., 2013). While CAMK2 has been extensively implicated in NMDAR-dependent LTP, CAMK4 is less well studied in this context (Lisman et al., 2012). CAMK4 has been shown to regulate Ca-dependent transcription via phosphorylation of various transcription factors including CREB (Enslen et al., 1995). PRKX is a cAMP-dependent protein kinase similar to PKA, a downstream target of DRD1A receptor activation which is a requirement for LTP (Li, 2011). It remains to be seen what role CAMK4 and PRKX might play in LTP in the context of LID and whether they are involved in the reported loss of depotentiation. In the indirect pathway, an upregulation of GUCY1A3 and downregulation of GNAO1 underscored the involvement of LTD. LID has been shown to be associated with a loss of corticostriatal LTD (Calabresi et al., 2000). Previous work has shown that this effect can be reversed by pharmacological treatment with phosphodiesterase inhibitors via modulation of cGMP levels. This is consistent with our finding of alterations in GUCY1A3, which encodes Guanylate cyclase soluble subunit alpha-3.

Interestingly, we also found that CCK (cholecystokinin) was upregulated in the direct versus indirect pathway in animals treated with both acute LDOPA or chronic LDOPA. Activation of the central CCK receptor (CCK-B) has been shown to augment LTP in guinea pig hippocampus (Yasui & Kawasaki, 1995). Furthermore, the CCK analogue CCK-8S has been shown to inhibit LID in parkinsonian squirrel monkeys (Boyce et al., 1990). While the striatum has been shown to have high concentrations of CCK and abundantly expresses CCK receptors, the role of CCK in dopaminergic regulation of LTP and its relationship with the mechanism by which it may inhibit LID has not been studied and thus represents a new avenue of research (Okonkwo & Adeyinka, 2019).

### Comparison to previous studies in DRD1A and DRD2 EGFP-L10a expressing BacTRAP animals

In 2014, Heiman and colleagues performed a study in DRD1A and DRD2 EGFP-L10a expressing BacTRAP animals, exploring alterations in gene expression resulting from dopamine denervation and in response to treatment with two different doses of LDOPA for 9 days (Heiman et al., 2014). This study noted only a small number of gene expression changes in the indirect pathway in response to LDOPA, whereas the direct pathway exhibited profound alterations in CREB, AP-1 and ERK-mediated signalling, with many of the associated genes correlating with the dose of LDOPA. Although differences in the experimental paradigms employed in the present study and the work of Heiman et al., preclude the direct comparison of the datasets, importantly, there is some overlap in the pathways identified, notably alterations in MAPK signalling in the direct pathway. Our study is distinct from the Heiman et al., study in two critical ways. First, Heiman et al., focussed on genes implicated with dopamine depletion and the *severity* of LID and investigated the effects of chronic treatment with high dose LDOPA vs low dose LDOPA, whereas our study examined the effects of *chronic* LDOPA treatment vs *acute* LDOPA (single dose). The rationale for our treatment regimen was to identify changes in gene expression resulting from repeated vs initial exposure to LDOPA as these genes might best reflect pathological processes implicated in the development of LID in response to repeated exposure. Second, we examined tissues collected 40 minutes after the last dose of LDOPA, whereas Heiman et al., examined tissues 3 hours and 20 minutes after the last LDOPA dose. Although we have not performed a pharmacokinetic analysis in the present study, the use of LDOPA in 6-OHDA lesioned mice is extensively described in the literature and it is well established that LDOPA reaches peak plasma concentrations 30-60 minutes post administration (Cenci & Crossman, 2018; Eriksson et al., 1984; Lundblad et al., 2004; Putterman et al., 2007; Winkler et al., 2002). After this point, LDOPA declines in its bioavailability and returns to control values approximately 120 minutes post administration (Kääriäinen et al., 2012; Peng et al., 2019; Spencer & Wooten, 1984). Thus, in contrast to Heiman and colleagues, our study is designed to capture gene expression concurrent with the expression of LID. Accordingly, there are meaningful differences between the two studies that have a clear impact on the hypothesis being tested and are reflected in the different genes and pathways identified in the two pieces of work.

### Potential caveats

The present work needs to be considered in light of some caveats. First, our data are restricted to those that manifest in alteration of RNA expression levels and therefore do not detect LID-related processes that involve enzymatic reactions (e.g. phosphorylation) or protein-protein interactions. For example, mTOR signalling has been shown to be a downstream target in the MAPK pathway associated with LID, however as this observation is associated with signalling events, not a change in expression *per se,* this was not detected in our study (Jin et al., 2010; Zhu et al., 2020). Second, a recent study reported that RNAseq increased the sensitivity of the TRAP approach by ∼10-fold compared to the microarray approach employed here, suggesting that had TRAP-RNAseq been used additional genes of interest may have been found (Montalban et al., 2022). Indeed, the TRAP-RNAseq model has led to the discovery of previously unknown distinct MSN subpopulations involved in locomotor control (Fieblinger, 2021). Third, while our study focused on changes in gene expression in the striatum, other studies have demonstrated that chronic LDOPA can induce changes in gene expression in other areas of the brain such as the frontal cortex, as well as in cell types other than MSNs, including immune and endothelial cells (Radlicka et al., 2021). Lastly, our study did not consider changes in DNA methylation which have been implicated in the development and maintenance of LID (Figge et al., 2016).

## Conclusion

In summary, we have employed a powerful experimental paradigm, combining BacTRAP (to selectively isolate RNA from either DRD1A expressing striatonigral or DRD2 expressing striatopallidal MSNs), with a widely used rodent model of LID, to reveal changes in translational gene expression following repeated LDOPA treatment. Our findings have revealed several novel translational changes implicated in LID. Our data highlight the involvement of several genes in the MAPK pathway, calcium signaling and LTP/LTD in maladaptive responses to chronic treatment with LDOPA. These findings have implicated the potential of targeting these genes and pathways as novel therapeutic interventions to prevent the development of LID, a critical unmet need in the treatment of PD.

---

**Supplementary Figure 1.**
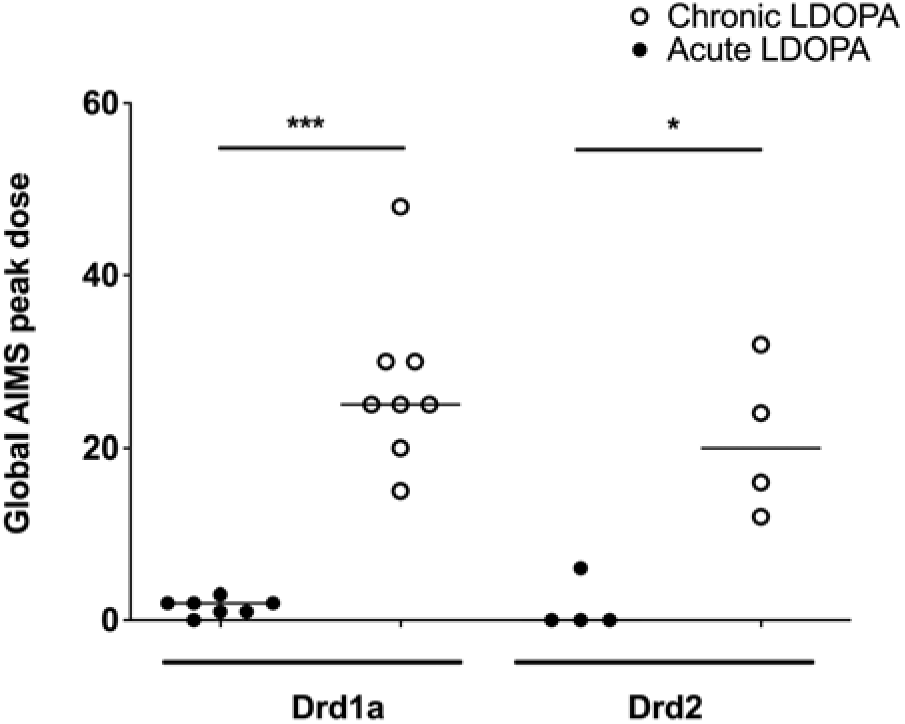
Abnormal involuntary movements (AIMs) in Drd1a and Drd2 EGFP-L10a expressing MSNs following acute or chronic treatment with LDOPA in a mouse model of PD. On day 22, AIMs were assessed in all animals. Data are presented from each individual animal expressed as the global AIMs score 40 minutes post treatment with LDOPA methyl ester/benserazide (6/1.5mg/kg). *=P<0.05, ***=P<0.001 Kruskal-Wallis test followed by Dunn’s Multiple Comparison Test.

**Supplementary Figure 2.**
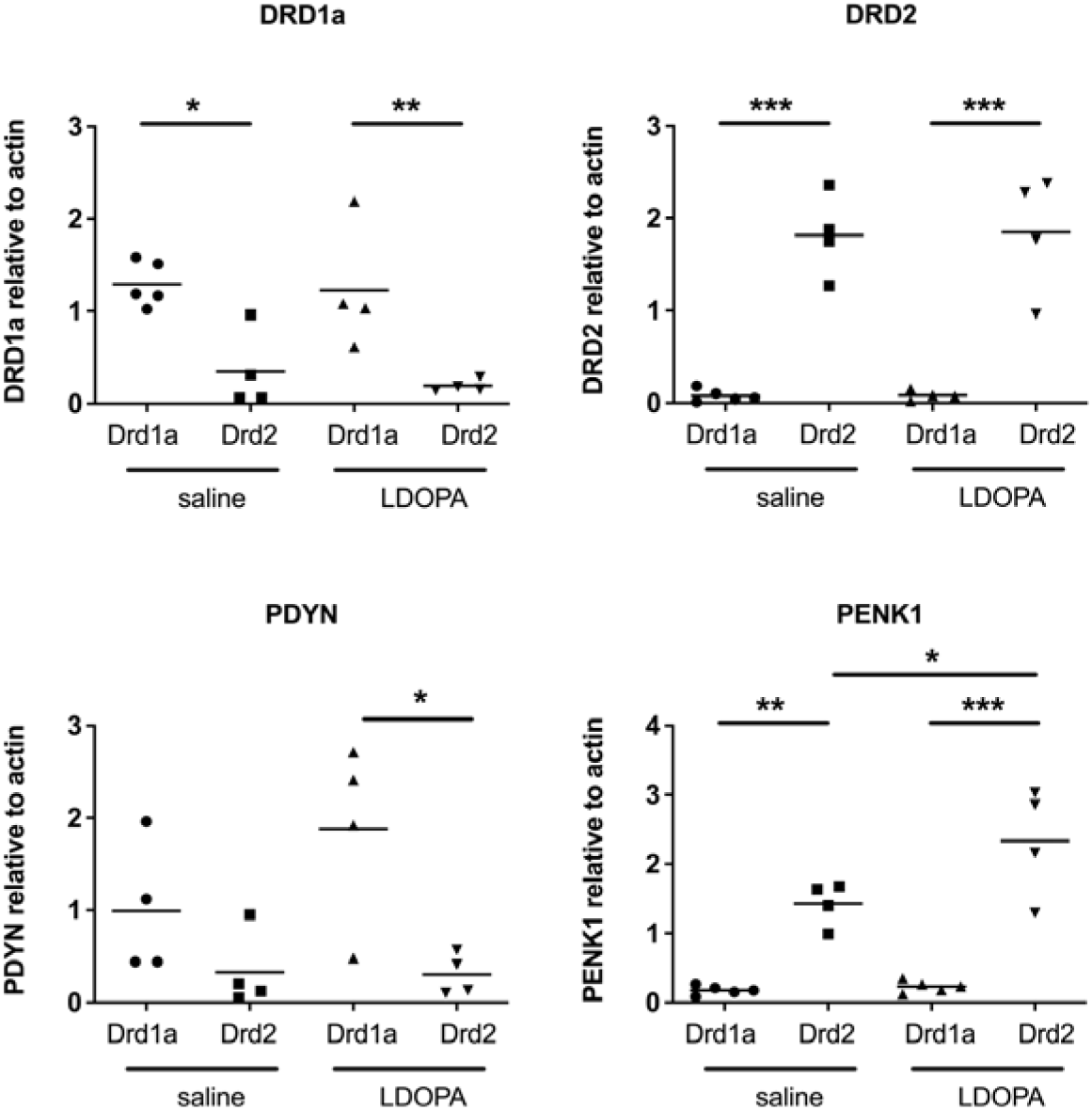
Q-RTPCR validation of known gene expression patterns in Drd1a and Drd2 EGFP-L10a expressing MSNs following acute or chronic treatment with LDOPA in a mouse model of PD. Individual values for each gene of interest are presented for each animal relative to actin. *=P<0.05, **=P<0.01, ***=P<0.001, One way ANOVA with Bonferroni’s Multiple Comparison Test.

**Supplementary Figure 3.**
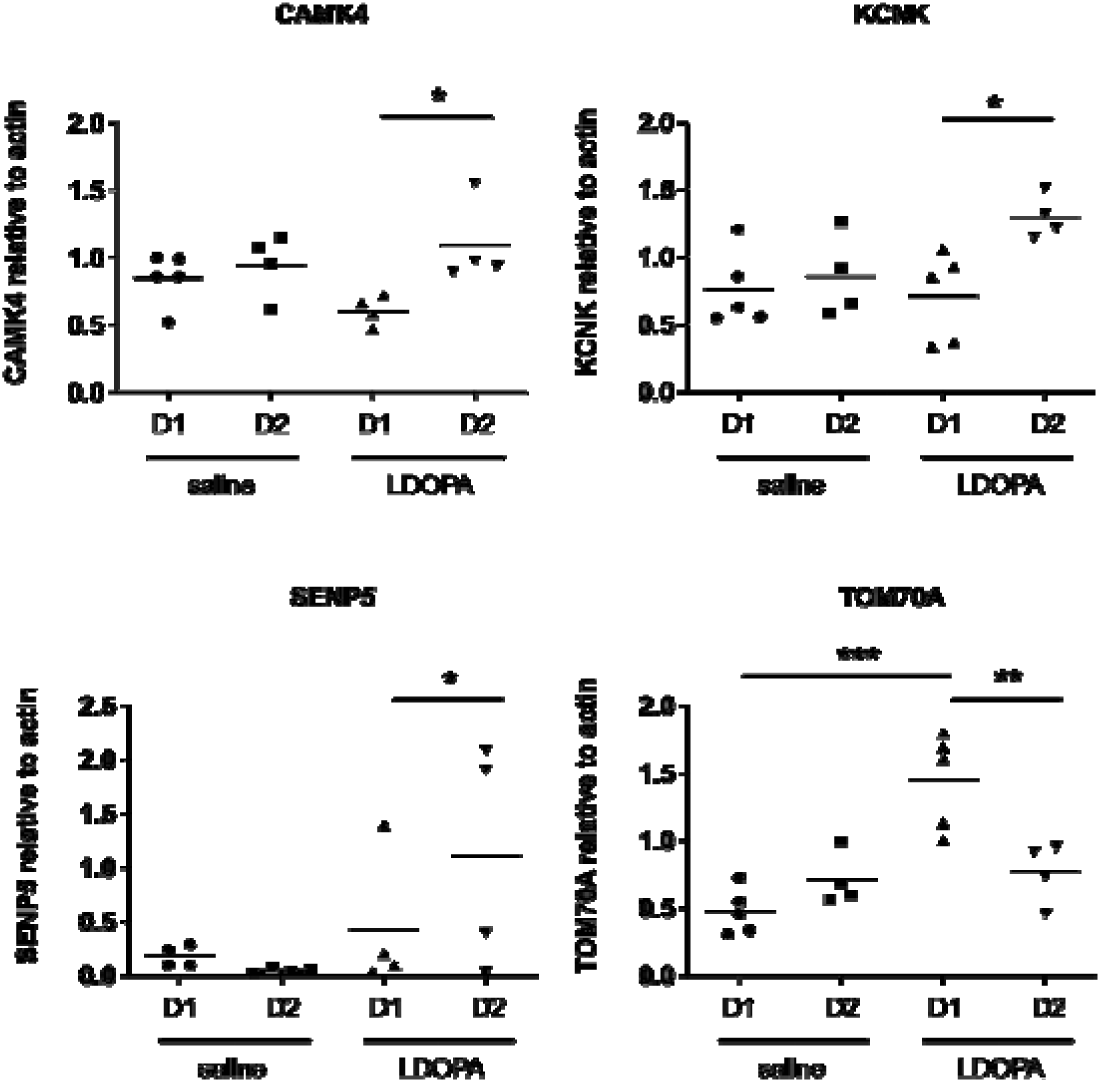
Q-RTPCR validation of gene expression patterns identified using microarray in Drd1a and Drd2 EGFP-L10a expressing MSNs following acute or chronic treatment with LDOPA in a mouse model of PD. Individual values for each gene of interest are presented for each animal relative to actin. *=P<0.05, **=P<0.01, ***=P<0.001, One way ANOVA with Bonferroni’s Multiple Comparison Test.

## Abbreviations

LDOPA: L-dihydroxyphenylalanine
TRAP: Translating ribosomal affinity purification
Bac: Bacterial artificial chromosome
MSNs: Medium spiny neurons
6-OHDA: 6-hydroxydopamine
PD: Parkinson’s disease
LID: LDOPA induced dyskinesia
AIMs: Abnormal involuntary movements
EGFP: Enhanced green fluorescent protein
i.p.: intraperitoneally
MFB: Median forebrain bundle
TH: Tyrosine hydroxylase
HSD: Honest significant difference
KEGG: Kyoto Encyclopedia of Genes and Genomes
FDR: False discovery rate
GO: Gene ontology
MAPK: Mitogen-activated protein kinase
LTP: Long term potentiation
LTD: Long term depression
PDYN: Prodynorphin
PENK1: Preproenkephalin 1
CCK: Cholecystokinin

## Declarations

### Ethics approval and consent to participate

This study was carried out in strict accordance with the standards set by the Canadian Council on Animal Care. The Animal Use Protocol was approved by the Animal Care Committee at the University of Toronto (approved protocol number 20008997).

### Consent for publication

Not applicable.

### Availability of data and materials

The datasets used and/or analysed during the current study are available from the corresponding author on request.

### Authors’ contributions

SJ edited the manuscript, read and approved the final manuscript.

MG performed data analysis and read and approved the final manuscript.

CV performed data analysis and r read and approved the final manuscript.

LN-H contributed to the conception and design of the experiment and read and approved the final manuscript.

NPV contributed to the conception and design of the experiment.

NV carried out the research, conducted the experiments, and performed data analysis.

NV drafted the first manuscript and created figures and tables.

NV revised the manuscript.

All authors read and approved the final manuscript.

## Acknowledgements

Not applicable

---

**Supplemental Table 1.** Behavioural description of each AIM subtype and rating. The AIMs scale was used to rate the animals from 1-4 based on the maximum severity of behaviour observed in each 1 min period for a span of 3 hours.

| Subtype | Rating | Behaviour |
| --- | --- | --- |
| All Limb | 0 | normal motor repertoire |
|  | 1 | paw displaced repetitively to and from snout or cage floor |
|  | 2 | visible displacement of whole limb either sideways or up/down |
|  | 3 | large displacement of whole limb with contraction of shoulder muscles |
|  | 4 | vigorous limb displacement of maximum amplitude |
| Axial | 1 | sustained deviation head & neck ~30° |
|  | 2 | sustained deviation head & neck 30° - 60° |
|  | 3 | sustained twisting head, neck and upper trunk 60° - 90° |
|  | 4 | sustained twisting head, neck and upper trunk > 90° with loss of balance |
| Orolingual | 1 | twitching facial muscles, small masticatory movements no jaw opening |
|  | 2 | twitching facial muscles, noticeable mastication, occasional jaw opening |
|  | 3 | frequent jaw opening, occasional tongue protrusion |
|  | 4 | maximal facial and masticatory movements, jaw opening and tongue protrusion |

**Supplemental Table 2.**
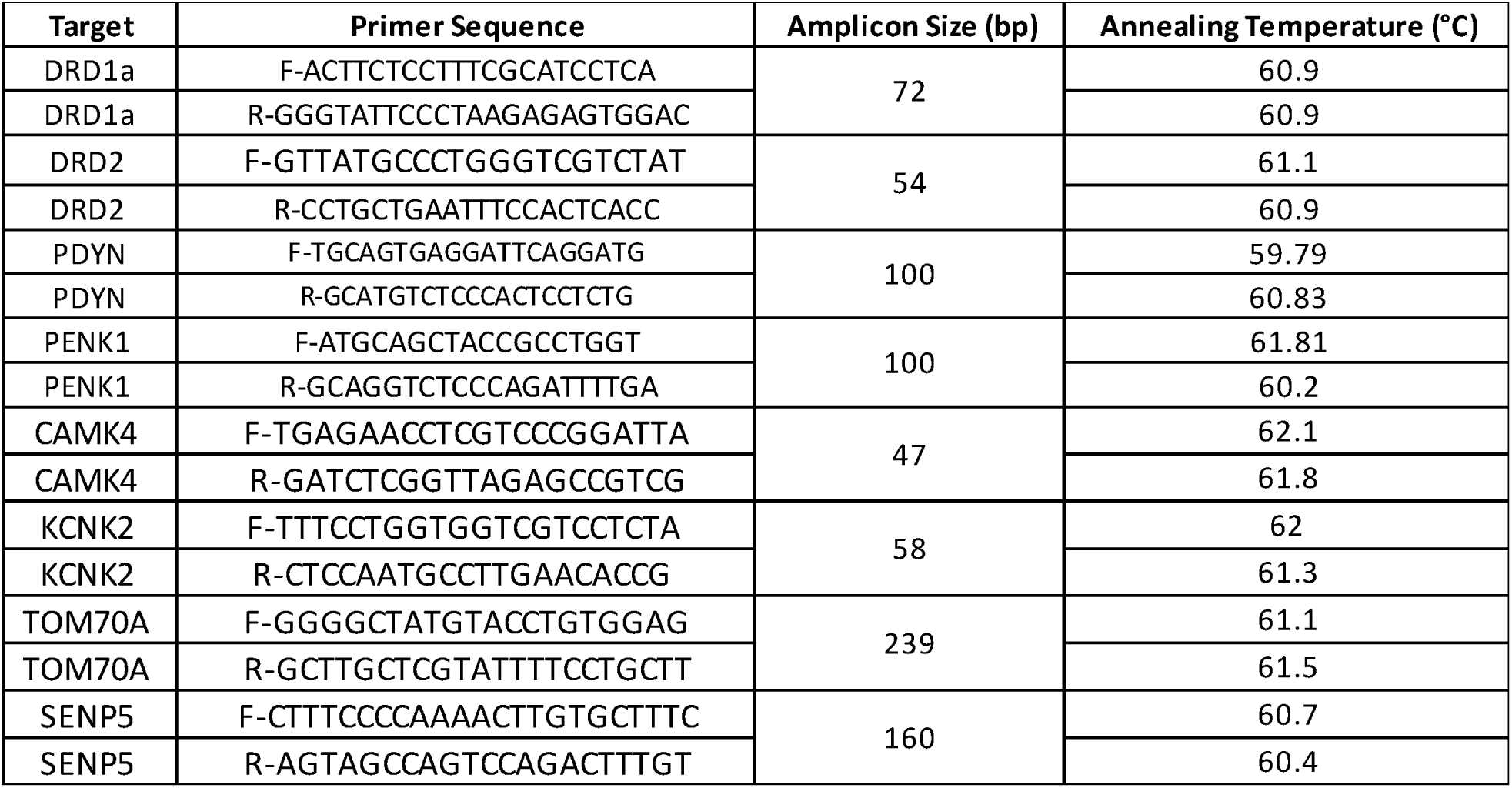
qRT-PCR primer sequences. Complete list of all primers used in the study.

**Supplemental Table 3.** Confirmation of lesion of the nigrostriatal pathway. Animals’ rotational behaviour was assessed for 10 min and the number of ipsiversive rotations calculated as a percentage of the total number of rotations in either direction. The number of nigral tyrosine hydroxylase (TH) immunoreactive cells was assessed in 3 consecutive sections from each animal and the number of cells in the lesioned hemisphere calculated as a percentage of those in the intact hemisphere . Data are presented as the mean +/-standard error of the mean per group.

| Genotype | Treatment | Number of animals | % Ipsiversive rotation | % TH cell loss |
| --- | --- | --- | --- | --- |
| DRD1-EGFP | Saline | 7 | 97 ± 2 | 93 ± 3 |
| DRD1A-EGFP | L-DOPA | 8 | 95 ± 1 | 91 ± 3 |
| DRD2-EGFP | Saline | 4 | 95 ± 1 | 91 ± 2 |
| DRD2-EGFP | L-DOPA | 4 | 100 ± 0 | 90 ± 3 |

## References

Albin, R. L., Young, A. B., & Penney, J. B. (1989). The functional anatomy of basal ganglia disorders. Trends in Neurosciences, 12(10), 366–375. 10.1016/0166-2236(89)90074-X

Alexander, G. E., & Crutcher, M. D. (1990). Functional architecture of basal ganglia circuits: neural substrates of parallel processing. Trends in Neurosciences, 13(7), 266–271. 10.1016/0166-2236(90)90107-L

AlShimemeri, S., Fox, S. H., & Visanji, N. P. (2020). Emerging drugs for the treatment of L-DOPA-induced dyskinesia: an update. Expert Opinion on Emerging Drugs, 25(2), 131–144. 10.1080/14728214.2020.1763954

Anderson, K. A., & Kane, C. D. (1998). Ca2+/calmodulin-dependent protein kinase IV and calcium signaling. BioMetals, 11(4), 331–343. 10.1023/A:1009276932076

Angela Cenci, M., & Lundblad, M. (2007). Ratings of l-dopa-induced dyskinesia in the unilateral 6-ohda lesion model of parkinson’s disease in rats and mice. Current Protocols in Neuroscience, 41(1), 1–23. 10.1002/0471142301.ns0925s41

Asrar, S., Meng, Y., Zhou, Z., Todorovski, Z., Huang, W. W., & Jia, Z. (2009). Regulation of hippocampal long-term potentiation by p21-activated protein kinase 1 (PAK1). Neuropharmacology, 56(1), 73–80. 10.1016/j.neuropharm.2008.06.055

Azkona, G., Sagarduy, A., Aristieta, A., Vazquez, N., Zubillaga, V., Ruíz-Ortega, J. A., Pérez-Navarro, E., Ugedo, L., & Sánchez-Pernaute, R. (2014). Buspirone anti-dyskinetic effect is correlated with temporal normalization of dysregulated striatal DRD1 signalling in l-DOPA-treated rats. Neuropharmacology, 79, 726–737. 10.1016/j.neuropharm.2013.11.024

Balliet, A. G., Hollander, M. C., Fornace, A. J., Hoffman, B., & Liebermann, D. A. (2003). Comparative analysis of the genetic structure and chromosomal mapping of the murine Gadd45g/CR6 gene. DNA and Cell Biology, 22(7), 457–468. 10.1089/104454903322247334

Beck, G., Singh, A., Zhang, J., Potts, L. F., Woo, J. M., Park, E. S., Mochizuki, H., Maral Mouradian, M., & Papa, S. M. (2019). Role of striatal ΔFosB in L-Dopa–induced dyskinesias of parkinsonian nonhuman primates. Proceedings of the National Academy of Sciences of the United States of America, 116(37), 18664–18672. 10.1073/pnas.1907810116

Bell, K. F. S., Bent, R. J., Meese-Tamuri, S., Ali, A., Forder, J. P., & Aarts, M. M. (2013). Calmodulin kinase IV-dependent CREB activation is required for neuroprotection via NMDA receptor-PSD95 disruption. Journal of Neurochemistry, 126(2), 274–287. 10.1111/jnc.12176

Bezard, E., Brotchie, J. M., & Gross, C. E. (2001). Pathophysiology of levodopa-induced dyskinesia: potential for new therapies. Nature Reviews Neuroscience, 2(8), 577–588. 10.1038/35086062

Borgkvist, A., Lieberman, O. J., & Sulzer, D. (2018). Synaptic plasticity may underlie L-DOPA induced dyskinesia. Current Opinion in Neurobiology, 48, 71–78. 10.1016/j.conb.2017.10.021

Boyce, S., Rupniak, N., Steventon, M., & Iversen, S. D. (1990). CCK-8S inhibits l-dopa-induced dyskinesias in parkinsonian squirrel monkeys. Neurology, 40(4), 717–718. 10.1212/wnl.40.4.717

Calabresi, P., Giacomini, P., Centonze, D., & Bernardi, G. (2000). Levodopa-induced dyskinesia: a pathological form of striatal synaptic plasticity? Annals of Neurology, *47*(4 Suppl 1), S60-8; discussion S68-9. http://www.ncbi.nlm.nih.gov/pubmed/10762133

Calabresi, P., Picconi, B., Tozzi, A., & Di Filippo, M. (2007). Dopamine-mediated regulation of corticostriatal synaptic plasticity. Trends in Neurosciences, 30(5), 211–219. 10.1016/J.TINS.2007.03.001

Calabresi, P., Picconi, B., Tozzi, A., Ghiglieri, V., & Di Filippo, M. (2014). Direct and indirect pathways of basal ganglia: A critical reappraisal. Nature Neuroscience, 17(8), 1022–1030. 10.1038/nn.3743

Cattabeni, F., Gardoni, F., & Di Luca, M. (1999). Pathophysiological implications of the structural organization of the excitatory synapse. European Journal of Pharmacology, 375(1–3), 339–347. 10.1016/S0014-2999(99)00299-X

Cenci, M. A., & Crossman, A. R. (2018). Animal models of l-dopa-induced dyskinesia in Parkinson’s disease. Movement Disorders, 33(6), 889–899. 10.1002/mds.27337

Cenci, M. A., Jörntell, H., & Petersson, P. (2018). On the neuronal circuitry mediating l-DOPA-induced dyskinesia. Journal of Neural Transmission, 125(8), 1157–1169. 10.1007/s00702-018-1886-0

Cenci, M. A., & Konradi, C. (2010). Maladaptive striatal plasticity in l-DOPA-induced dyskinesia. Progress in Brain Research, 183(C), 209–233. 10.1016/S0079-6123(10)83011-0

Chen, J., Abi-Daoud, M., Wang, A., Yang, X., Zhang, X., Feilotter, H. E., & Tron, V. A. (2013). Stathmin 1 is a potential novel oncogene in melanoma. Oncogene, 32(10), 1330–1337. 10.1038/onc.2012.141

Ciccarelli, A., & Giustetto, M. (2014). Role of ERK signaling in activity-dependent modifications of histone proteins. Neuropharmacology, 80, 34–44. 10.1016/j.neuropharm.2014.01.039

Costa, C., Sgobio, C., Siliquini, S., Tozzi, A., Tantucci, M., Ghiglieri, V., Di Filippo, M., Pendolino, V., de Iure, A., Marti, M., Morari, M., Spillantini, M. G., Latagliata, E. C., Pascucci, T., Puglisi-Allegra, S., Gardoni, F., Di Luca, M., Picconi, B., & Calabresi, P. (2012). Mechanisms underlying the impairment of hippocampal long-term potentiation and memory in experimental Parkinson’s disease. BrainfZ: A Journal of Neurology, 135(Pt 6), 1884–1899. 10.1093/brain/aws101

Deane, K. H. O., Flaherty, H., Daley, D. J., Pascoe, R., Penhale, B., Clarke, C. E., Sackley, C., & Storey, S. (2015). Priority setting partnership to identify the top 10 research priorities for the management of parkinson’s disease. BMJ Open, 4(12). 10.1136/bmjopen-2014-006434

Dodel, R. C., Berger, K., & Oertel, W. H. (2001). Health-related quality of life and healthcare utilisation in patients with Parkinson’s disease: Impact of motor fluctuations and dyskinesias. PharmacoEconomics, 19(10), 1013–1038. 10.2165/00019053-200119100-00004

Dougherty, J. D. (2017). The expanding toolkit of translating ribosome affinity purification. Journal of Neuroscience, 37(50), 12079–12087. 10.1523/JNEUROSCI.1929-17.2017

Doyle, J. P., Dougherty, J. D., Heiman, M., Schmidt, E. F., Stevens, T. R., Ma, G., Bupp, S., Shrestha, P., Shah, R. D., Doughty, M. L., Gong, S., Greengard, P., & Heintz, N. (2008). Application of a Translational Profiling Approach for the Comparative Analysis of CNS Cell Types. Cell, 135(4), 749–762. 10.1016/j.cell.2008.10.029

Dyavar, S. R., Potts, L. F., Beck, G., Dyavar Shetty, B. L., Lawson, B., Podany, A. T., Fletcher, C. V., Amara, R. R., & Papa, S. M. (2020). Transcriptomic approach predicts a major role for transforming growth factor beta type 1 pathway in L-Dopa-induced dyskinesia in parkinsonian rats. Genes, Brain and Behavior, 19(8). 10.1111/gbb.12690

Enslen, H., Tokumitsu, H., & Soderling, T. R. (1995). Phosphorylation of CREB by CaM-Kinase IV activated by CaM-kinase IV kinase. Biochemical and Biophysical Research Communications, 207(3), 1038–1043. 10.1006/bbrc.1995.1289

Eriksson, T., Magnusson, T., Carlsson, A., Linde, A., & Granérus, A. K. (1984). “On-off” phenomenon in Parkinson’s disease: Correlation to the concentration of dopa in plasma. Journal of Neural Transmission, 59(3), 229–240. 10.1007/BF01250010

Fabbrini, A., & Guerra, A. (2021). Pathophysiological Mechanisms and Experimental Pharmacotherapy for L-Dopa-Induced Dyskinesia. Journal of Experimental Pharmacology, 13, 469–485. 10.2147/JEP.S265282

Fabbrini, G., Brotchie, J. M., Grandas, F., Nomoto, M., & Goetz, C. G. (2007). Levodopa-induced dyskinesias. Movement Disorders, 22(10), 1379–1389. 10.1002/mds.21475

Feng, T., Xu, J., He, P., Chen, Y., Fang, R., & Shao, X. (2017). Decrease in stathmin expression by arsenic trioxide inhibits the proliferation and invasion of osteosarcoma cells via the MAPK signal pathway. Oncology Letters, 14(2), 1333–1340. 10.3892/ol.2017.6347

Fieblinger, T. (2021). Striatal Control of Movement: A Role for New Neuronal (Sub-) Populations? Frontiers in Human Neuroscience, 15, 697284. 10.3389/fnhum.2021.697284

Fieblinger, T., Graves, S. M., Sebel, L. E., Alcacer, C., Plotkin, J. L., Gertler, T. S., Chan, C. S., Heiman, M., Greengard, P., Cenci, M. A., & Surmeier, D. J. (2014). Cell type-specific plasticity of striatal projection neurons in parkinsonism and L-DOPA-induced dyskinesia. Nature Communications, 5. 10.1038/ncomms6316

Figge, D. A., Eskow Jaunarajs, K. L., & Standaert, D. G. (2016). Dynamic DNA methylation regulates levodopa-induced Dyskinesia. Journal of Neuroscience, 36(24), 6514–6524. 10.1523/JNEUROSCI.0683-16.2016

Fiorentini, C., Rizzetti, M. C., Busi, C., Bontempi, S., Collo, G., Spano, P. F., & Missale, C. (2006). Loss of synaptic D1 dopamine/N-methyl-D-aspartate glutamate receptor complexes in L-DOPA-induced dyskinesia in the rat. Molecular Pharmacology, 69(3), 805–812. 10.1124/mol.105.016667

Fisone, G., & Bezard, E. (2011). Molecular mechanisms of l-DOPA-induced dyskinesia. International Review of Neurobiology, 98, 95–122. 10.1016/B978-0-12-381328-2.00004-3

Fortin, D. A., Davare, M. A., Srivastava, T., Brady, J. D., Nygaard, S., Derkach, V. A., & Soderling, T. R. (2010). Long-term potentiation-dependent spine enlargement requires synaptic Ca2+-permeable AMPA receptors recruited by CaM-kinase I. Journal of Neuroscience, 30(35), 11565– 11575. 10.1523/JNEUROSCI.1746-10.2010

George Paxinos, & Keith B.J. Franklin. (2013). Paxinos and Franklin’s the mouse brain in stereotaxic coordinates (4th ed., Vol. 1). Elsevier Academic Press.

Girasole, A. E., Lum, M. Y., Nathaniel, D., Bair-Marshall, C. J., Guenthner, C. J., Luo, L., Kreitzer, A. C., & Nelson, A. B. (2018). A Subpopulation of Striatal Neurons Mediates Levodopa-Induced Dyskinesia. Neuron, 97(4), 787–795.e6. 10.1016/j.neuron.2018.01.017

Heiman, M., Kulicke, R., Fenster, R. J., Greengard, P., & Heintz, N. (2014). Cell type-specific mRNA purification by translating ribosome affinity purification (TRAP). Nature Protocols, 9(6), 1282– 1291. 10.1038/nprot.2014.085

Heiman, M., Schaefer, A., Gong, S., Peterson, J. D., Day, M., Ramsey, K. E., Suárez-Fariñas, M., Schwarz, C., Stephan, D. A., Surmeier, D. J., Greengard, P., & Heintz, N. (2008). A Translational Profiling Approach for the Molecular Characterization of CNS Cell Types. Cell, 135(4), 738–748. 10.1016/j.cell.2008.10.028

Huot, P., Johnston, T. H., Koprich, J. B., Fox, S. H., & Brotchie, J. M. (2013). The pharmacology of L-DOPA-induced dyskinesia in Parkinson’s disease. Pharmacological Reviews, 65(1), 171–222. 10.1124/pr.111.005678

Hutny, M., Hofman, J., Klimkowicz-Mrowiec, A., & Gorzkowska, A. (2021). Current Knowledge on the Background, Pathophysiology and Treatment of Levodopa-Induced Dyskinesia-Literature Review. Journal of Clinical Medicine, 10(19). 10.3390/jcm10194377

Jenner, P. (2008). Molecular mechanisms of L-DOPA-induced dyskinesia. Nature Reviews Neuroscience, 9(9), 665–677. 10.1038/nrn2471

Jin, C. M., Yang, Y. J., Huang, H. S., Kai, M., & Lee, M. K. (2010). Mechanisms of L-DOPA-induced cytotoxicity in rat adrenal pheochromocytoma cells: Implication of oxidative stress-related kinases and cyclic AMP. Neuroscience, 170(2), 390–398. 10.1016/j.neuroscience.2010.07.039

Kääriäinen, T. M., Käenmäki, M., Forsberg, M. M., Oinas, N., Tammimäki, A., & Männistö, P. T. (2012). Unpredictable rotational responses to L-dopa in the rat model of Parkinson’s disease: The role of L-dopa pharmacokinetics and striatal dopamine depletion. Basic and Clinical Pharmacology and Toxicology, 110(2), 162–170. 10.1111/j.1742-7843.2011.00782.x

Koga, Y., Tsurumaki, H., Aoki-Saito, H., Sato, M., Yatomi, M., Takehara, K., & Hisada, T. (2019). Roles of cyclic AMP response element binding activation in the ERK1/2 and p38 MAPK signalling pathway in central nervous system, cardiovascular system, osteoclast differentiation and mucin and cytokine production. International Journal of Molecular Sciences, 20(6). 10.3390/ijms20061346

Li, X. (2011). Phosphorylation, protein kinases and ADPKD. Biochimica et Biophysica Acta - Molecular Basis of Disease, 1812(10), 1219–1224. 10.1016/j.bbadis.2011.03.001

Lisman, J., Yasuda, R., & Raghavachari, S. (2012). Mechanisms of CaMKII action in long-term potentiation. Nature Reviews Neuroscience, 13(3), 169–182. 10.1038/nrn3192

Lundblad, M., Andersson, M., Winkler, C., Kirik, D., Wierup, N., & Cenci Nilsson, M. A. (2002). Pharmacological validation of behavioural measures of akinesia and dyskinesia in a rat model of Parkinson’s disease. European Journal of Neuroscience, 15(1), 120–132. 10.1046/j.0953-816x.2001.01843.x

Lundblad, M., Picconi, B., Lindgren, H., & Cenci, M. A. (2004). A model of L-DOPA-induced dyskinesia in 6-hydroxydopamine lesioned mice: Relation to motor and cellular parameters of nigrostriatal function. Neurobiology of Disease, 16(1), 110–123. 10.1016/j.nbd.2004.01.007

Lundblad, M., Usiello, A., Carta, M., Håkansson, K., Fisone, G., & Cenci, M. A. (2005). Pharmacological validation of a mouse model of L-DOPA-induced dyskinesia. Experimental Neurology, 194(1), 66–75. 10.1016/j.expneurol.2005.02.002

Maxwell, M. A., & Muscat, G. E. O. (2006). The NR4A subgroup: immediate early response genes with pleiotropic physiological roles. Nuclear Receptor Signaling, 4(1), e002. 10.1621/nrs.04002

Montalban, E., Giralt, A., Taing, L., Schut, E. H. S., Supiot, L. F., Castell, L., Nakamura, Y., de Pins, B., Pelosi, A., Goutebroze, L., Tuduri, P., Wang, W., Neiburga, K. D., Vestito, L., Castel, J., Luquet, S., Nairn, A. C., Hervé, D., Heintz, N., … Girault, J. A. (2022). Translational profiling of mouse dopaminoceptive neurons reveals region-specific gene expression, exon usage, and striatal prostaglandin E2 modulatory effects. Molecular Psychiatry, 27(4), 2068–2079. 10.1038/s41380-022-01439-4

Mosharov, E. V., Borgkvist, A., & Sulzer, D. (2015). Presynaptic effects of levodopa and their possible role in dyskinesia. Movement Disorders, 30(1), 45–53. 10.1002/mds.26103

Obeso, J. A., Olanow, C. W., & Nutt, J. G. (2000). Levodopa motor complications in Parkinson’s disease. Trends in Neurosciences, *23*(10 SUPPL.). 10.1016/S1471-1931(00)00031-8

Okonkwo, O., & Adeyinka, A. (2019). Biochemistry, Cholecystokinin (CCK). StatPearls. http://www.ncbi.nlm.nih.gov/pubmed/30480943

Pagliaroli, L., Widomska, J., Nespoli, E., Hildebrandt, T., Barta, C., Glennon, J., Hengerer, B., & Poelmans, G. (2019). Riluzole Attenuates L-DOPA-Induced Abnormal Involuntary Movements Through Decreasing CREB1 Activity: Insights from a Rat Model. Molecular Neurobiology, 56(7), 5111–5121. 10.1007/s12035-018-1433-x

Peng, Q., Zhong, S., Tan, Y., Zeng, W., Wang, J., Cheng, C., Yang, X., Wu, Y., Cao, X., & Xu, Y. (2019). The Rodent Models of Dyskinesia and Their Behavioral Assessment. Frontiers in Neurology, 10, 1016. 10.3389/fneur.2019.01016

Pérez-Sen, R., Queipo, M. J., Gil-Redondo, J. C., Ortega, F., Gómez-Villafuertes, R., Miras-Portugal, M. T., & Delicado, E. G. (2019). Dual-specificity phosphatase regulation in neurons and glial cells. International Journal of Molecular Sciences, 20(8). 10.3390/ijms20081999

Picconi, B., Piccoli, G., & Calabresi, P. (2012). Synaptic dysfunction in Parkinson’s disease. Advances in Experimental Medicine and Biology, 970, 553–572. 10.1007/978-3-7091-0932-8_24

Putterman, D. B., Munhall, A. C., Kozell, L. B., Belknap, J. K., & Johnson, S. W. (2007). Evaluation of levodopa dose and magnitude of dopamine depletion as risk factors for levodopa-induced dyskinesia in a rat model of Parkinson’s disease. Journal of Pharmacology and Experimental Therapeutics, 323(1), 277–284. 10.1124/jpet.107.126219

Radlicka, A., Kamińska, K., Borczyk, M., Piechota, M., Korostyński, M., Pera, J., Lorenc-Koci, E., & Parkitna, J. R. (2021). Effects of l-dopa on gene expression in the frontal cortex of rats with unilateral lesions of midbrain dopaminergic neurons. ENeuro, 8(1), 1–13. 10.1523/ENEURO.0234-20.2020

Rascol, O. (2000). Medical treatment of levodopa-induced dyskinesias. Annals of Neurology, *47*(4 Suppl 1), S179-88. http://www.ncbi.nlm.nih.gov/pubmed/10762146

Reyskens, K. M. S. E., & Arthur, J. S. C. (2016). Emerging Roles of the Mitogen and Stress Activated Kinases MSK1 and MSK2. Frontiers in Cell and Developmental Biology, 4(JUN), 56. 10.3389/fcell.2016.00056

Ryan, M. B., Bair-Marshall, C., & Nelson, A. B. (2018). Aberrant Striatal Activity in Parkinsonism and Levodopa-Induced Dyskinesia. Cell Reports, 23(12), 3438–3446.e5. 10.1016/j.celrep.2018.05.059

Santini, E., Alcacer, C., Cacciatore, S., Heiman, M., Hervé, D., Greengard, P., Girault, J. A., Valjent, E., & Fisone, G. (2009). L-DOPA activates ERK signaling and phosphorylates histone H3 in the striatonigral medium spiny neurons of hemiparkinsonian mice. Journal of Neurochemistry, 108(3), 621–633. 10.1111/J.1471-4159.2008.05831.X

Shrestha, Y., Schafer, E. J., Boehm, J. S., Thomas, S. R., He, F., Du, J., Wang, S., Barretina, J., Weir, B. A., Zhao, J. J., Polyak, K., Golub, T. R., Beroukhim, R., & Hahn, W. C. (2012). PAK1 is a breast cancer oncogene that coordinately activates MAPK and MET signaling. Oncogene, 31(29), 3397–3408. 10.1038/onc.2011.515

Spencer, S. E., & Wooten, G. F. (1984). Altered pharmacokinetics of L-dopa metabolism in rat striatum deprived of dopaminergic innervation. Neurology, 34(8), 1105–1108. 10.1212/wnl.34.8.1105

Spigolon, G., & Fisone, G. (2018). Signal transduction in l-DOPA-induced dyskinesia: from receptor sensitization to abnormal gene expression. Journal of Neural Transmission, 125(8), 1171–1186. 10.1007/s00702-018-1847-7

Steece-Collier, K., Collier, T. J., Lipton, J. W., Stancati, J. A., Winn, M. E., Cole-Strauss, A., Sellnow, R., Conti, M. M., Mercado, N. M., Nillni, E. A., Sortwell, C. E., Manfredsson, F. P., & Bishop, C. (2020). Striatal Nurr1, but not FosB expression links a levodopa-induced dyskinesia phenotype to genotype in Fisher 344 vs. Lewis hemiparkinsonian rats. Experimental Neurology, 330. 10.1016/j.expneurol.2020.113327

Surmeier, D. J., Graves, S. M., & Shen, W. (2014). Dopaminergic modulation of striatal networks in health and Parkinson’s disease. Current Opinion in Neurobiology, 29, 109–117. 10.1016/j.conb.2014.07.008

Visanji, N. P., Kamali Sarvestani, I., Creed, M. C., Shams Shoaei, Z., Nobrega, J. N., Hamani, C., & Hazrati, L.-N. (2015). Deep brain stimulation of the subthalamic nucleus preferentially alters the translational profile of striatopallidal neurons in an animal model of Parkinson’s disease. Frontiers in Cellular Neuroscience, 9(June), 221. 10.3389/fncel.2015.00221

Wancket, L. M., Frazier, W. J., & Liu, Y. (2012). Mitogen-activated protein kinase phosphatase (MKP)-1 in immunology, physiology, and disease. Life Sciences, 90(7–8), 237–248. 10.1016/j.lfs.2011.11.017

Winkler, C., Kirik, D., Björklund, A., & Cenci, M. A. (2002). L-DOPA-induced dyskinesia in the intrastriatal 6-hydroxydopamine model of Parkinson’s disease: Relation to motor and cellular parameters of nigrostriatal function. Neurobiology of Disease, 10(2), 165–186. 10.1006/nbdi.2002.0499

Yasui, M., & Kawasaki, K. (1995). CCKB-Receptor Activation Augments the Long-Term Potentiation in Guinea Pig Hippocampal Slices. The Japanese Journal of Pharmacology, 68(4), 441–447. 10.1254/jjp.68.441

Yokoyama, K., Hiyama, A., Arai, F., Nukaga, T., Sakai, D., & Mochida, J. (2013). C-Fos regulation by the MAPK and PKC pathways in intervertebral disc cells. PloS One, 8(9). 10.1371/journal.pone.0073210

Zhu, J., Xia, R., Liu, Z., Shen, J., Gong, X., Hu, Y., Chen, H., Yu, Y., Gao, W., Wang, C., & Wang, S. L. (2020). Fenvalerate triggers Parkinson-like symptom during zebrafish development through initiation of autophagy and p38 MAPK/mTOR signaling pathway. Chemosphere, 243. 10.1016/j.chemosphere.2019.125336

